# Primate innate immune responses to bacterial and viral pathogens reveals an evolutionary trade-off between strength and specificity

**DOI:** 10.1101/2020.07.23.218339

**Authors:** Mohamed Bayoumi Fahmy Hawash, Joaquin Sanz-Remón, Jean-Christophe Grenier, Jordan Kohn, Vania Yotova, Zach Johnson, Robert E. Lanford, Jessica F. Brinkworth, Luis B. Barreiro

**Affiliations:** CHU Sainte-Justine, University of Montreal, Montreal, Canada; Departamento de Fiscia Teórica, Universidad de Zaragoza, Zaragoza, Spain; Institute BIFI for Biocomputation and Physics of Complex Systems, Universidad de Zaragoza, Zaragoza, Spain; Montreal Heart Institute, University of Montreal, Montreal, Canada; Department of Neuroscience, Emory University, University of California San Diego, United States; Department of Psychiatry, College of Health Sciences, University of California San Diego, United States; Illumina, San Diego CA, United States; Southwest National Primate Research Center, Texas Biomedical Research Institute, San Antonio, TX 78227, USA; Department of Anthropology, University of Illinois Urbana-Champaign, United States; Carl R. Woese, Institute for Genomic Biology, University of Illinois Urbana-Champaign, United States; Department of Genetic Medicine, University of Chicago, Chicago, United States

**Keywords:** Pathogen-associated molecular patterns, primate evolution, early immune responses to infection, immunodeficiency viruses, Gram-negative bacteria

## Abstract

Despite their close genetic relatedness, apes and African and Asian monkeys (AAMs), strongly differ in their susceptibility to severe bacterial and viral infections that are important causes of human disease. Such differences between humans and other primates are thought to be a result, at least in part, of inter-species differences in immune response to infection. However, due to the lack of comparative functional data across species, it remains unclear in what ways the immune systems of humans and other primates differ. Here, we report the whole genome transcriptomic responses of ape species (human, common chimpanzee) and AAMs (rhesus macaque and olive baboon) to bacterial and viral stimulation. We find stark differences in the responsiveness of these groups, with apes mounting a markedly stronger early transcriptional response to both viral and bacterial stimulation, altering the transcription of ∼40% more genes than AAMs. Additionally, we find that genes involved in the regulation of inflammatory and interferon responses show the most divergent early transcriptional responses across primates and that this divergence is attenuated over time. Finally, we find that relative to AAMs, apes engage a much less specific immune response to different classes of pathogens during the early hours of infection, upregulating genes typical of anti-viral and anti-bacterial responses regardless of the nature of the stimulus. Overall, these findings suggest apes exhibit increased sensitivity to bacterial and viral immune stimulation, activating a broader array of defense molecules that may be beneficial for early pathogen killing at the potential cost of increased energy expenditure and tissue damage.

## INTRODUCTION

Despite being close evolutionary relatives, humans, chimpanzees and African and Asian monkeys exhibit inter-species differences in sensitivity to and manifestation of certain bacterial and viral pathogens that are major causes of mortality in humans (e.g. HIV/AIDS, Hepatitis C Virus, broad range of commensal Gram-negative bacteria commonly implicated in sepsis).^1-5^ Humans, for example, are highly sensitive to stimulation by the Gram-negative bacterial cell wall component hexa-acylated lipopolysaccharide (LPS), miniscule amounts of which (2-4ng/kg) can provoke inflammation, malaise and fever, and a slightly higher dose, septic shock (15 ug/kg).^1,6,7^ In contrast, baboons and macaques require doses nearly 10 fold higher in concentration to trigger similar symptoms.^5,8,9^ Pattern recognition receptors (PRRs) such as Toll-like receptors (TLRs) play a central role in the mediation of innate immune responses to pathogens^10^. The limited number of studies comparing leukocyte function after stimulation with TLR-detected pathogen-associated molecules (PAMPs) suggest that the differences in infectious diseases susceptibility noted between apes and AAMs is, in part, the outcome of lineage-specific evolution of early innate immune system regulation and signaling.^11-13^ Indeed, innate immune components responsible for detecting pathogens, including TLRs that sense Gram-negative bacteria and single-stranded RNA viruses, have been found to be under positive selection in primates.^14,15^

Despite stark differences in the manifestation of severe infections between apes and African and Asian monkeys (AAMs), there are few reports directly comparing the gene expression response across species to bacterial and viral pathogens.^11,12^ Further, previous studies relied mainly on isolated cell types to characterize immune responses across primates^11,16^, which does not faithfully reflect the nature of the innate immune response that is a product of the interaction between several cell populations^17^. To better understand the evolution of the primate immune system, this study compares the early responses of apes (humans and common chimpanzees) and AAMs (rhesus macaques and olive baboons) to bacterial and viral stimulants. Here, we report on the whole genome expression of total blood leukocytes from these four primate species responding to bacterial and viral stimulation during the first 24 hours of challenge. Our results show that apes and AAMs have diverged in sensitivity to specific microbial assaults, such that ape leukocyte responses favor robust antimicrobial power over pathogen specificity at the potential cost of increased energetic expenditure and bystander tissue damage.

## Results

### Evolutionary relationships explain most of the transcriptional response variation in primates to bacteria or viral stimulation

To assess differences in innate immune function between higher order primates in as close an approximation to *in vivo* as possible, we challenged whole blood from humans (*Homo sapiens*; N=6), common chimpanzees (*Pan troglodytes*; N=6), rhesus macaques (*Macaca mulattta*; N=6) and olive baboons (*Papio anubis*; N=8) with bacterial or viral *stimuli* via venous draw directly into a media culture tube containing either lipopolysaccharide (LPS) from *Escherichia coli* O111:B4, gardiquimod (GARD), a single-stranded RNA viral mimetic, or endotoxin-free water, as a negative control (Control). Blood was stimulated for 4 and 24 hours before the total leukocytes were isolated, and RNA extracted for RNA-sequencing (**Figure 1A**). We chose these two molecules because in mammals they are broad signals of infection by pathogen types for which there are well established differences in disease manifestation between apes and AAMs (e.g., immunodeficiency viruses, Hepatitis C, common commensal bacteria that cause Gram-negative bacterial sepsis).^1,3,4^ Following quality control filtering, we analyzed 151 high-quality RNA-sequencing profiles across species and treatment combinations (see Methods, **Table S1**). We focus our comparative analyses on the expression levels of 14,140 one-to-one (1:1) orthologous genes, taking into account potential biases in expression estimates due to differences in mappability between species (Methods).

**Figure 1.**
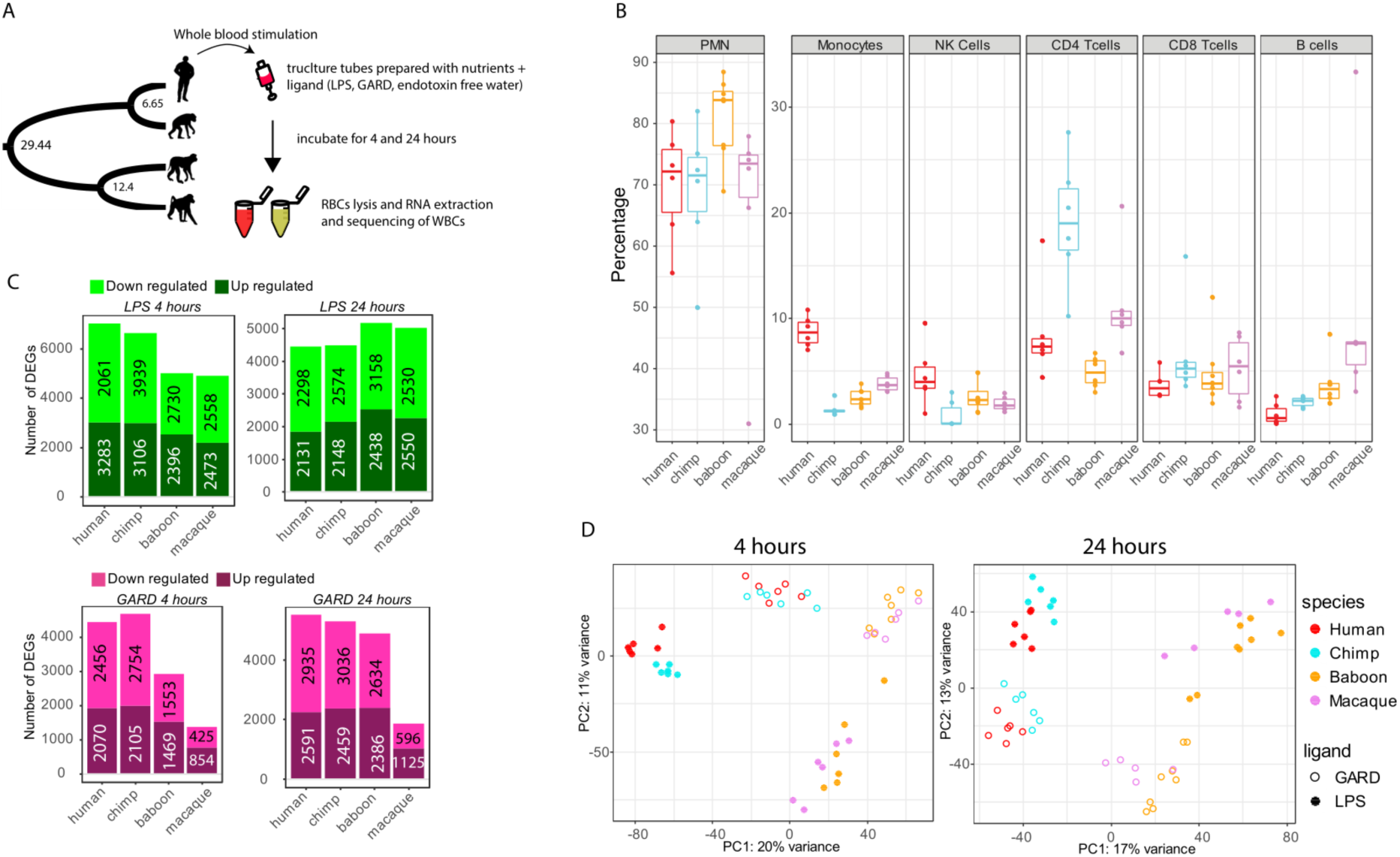
Characterizing innate immune response upon viral and bacterial stimulation of primate’s white blood cells. **(A)** Schematic representation of the study design. Whole blood samples from humans, common chimpanzees, rhesus macaques and olive baboons were stimulated with bacterial or viral *stimuli* via venous draw directly into a media culture tube containing either lipopolysaccharide (LPS), single stranded RNA viral mimetic gardiquimod (GARD), or endotoxin-free water, as a negative control (Control). At 4- and 24-hours post-stimulation white blood cells were isolated, and RNA extracted for RNA-sequencing. **(B)** Cell proportion of 6 populations of innate immune cells for all species. Species are indicated on x axis and proportions this population from total leukocytes is on y axis. Abbreviations are PMNs for polymorphonuclear cells and Natural killer for NK cells **(C)** Number of differentially expressed genes (DEGs; FDR<0.05) in response to LPS (top panels) and GARD (bottom panels) in each of the species at 4- and 24-hours post-stimulation The exact number of up- and down-regulated genes in each condition in each species are indicated on the bar charts. **(D)** Principal component analyses (PCA) performed on the log2 fold-change responses observed at 4 hours post LPS and GARD stimulation. PC1 primarily separates apes (human and chimpanzee) from AAM (macaque and baboon), and PC2 captures differences in immune response to bacterial or viral stimulation.

As whole blood contains a variety of leukocyte cell subtypes, we first characterized differences in total blood leukocyte composition between species using fluorescence-activated cell sorting (FACS). Leukocyte composition differs between species for all major subtypes measured, with the most notable differences an increase in the proportion of monocytes in humans (CD14^+^, *P* < 0.003) and helper T cells in chimpanzees (CD3^+^, CD4^+^; *P* = 0.0006 to 0.065), relative to other primates (**Figure 1B, Table S2**). Using linear models that account for variation in cell composition, we next identified genes that respond to LPS and GARD in each of the species, at each of the time points (see Methods). In all species, both treatments led to the up- or down-regulation of hundreds to thousands of genes (FDR<0.05, **Figure 1C, Table S3**). As expected, the transcriptional response to either stimulus was highly concordant across primates (Spearman’s r range 0.5 to 0.87 across all pairwise comparisons; **Figure S1**), with stronger correlations between closely related primates than between more distantly-related pairs of species (e.g., at LPS 4 hours Spearman’s r human vs chimpanzee = 0.84, human vs baboon = 0.50). Consistently, the first principal component (PC) of the log2 fold-change responses to both LPS and GARD accounted for ∼20% of the total variance in our dataset and separated apes (human and chimpanzee) from AAM (macaque and baboon) (t-test; *P* < 1×10^−10^ for both 4h and 24h, **Figure 1D**). The second PC captured differences in immune response to bacterial or viral stimulation (t-test; *P* < 1×10^−8^ for 4 and 24h; **Figure 1D**). We identified a set of 648 and 257 genes that early after stimulation (4 hours) showed a consistently strong response across all species to LPS or GARD, respectively (defined as genes with |log2 FC| > 1 and FDR < 0.05 in all species, **Table S4**). These genes include most of the key transcription factors involved in the regulation of innate immune responses to bacterial (e.g., *NFKB1*/*2*) and viral pathogens (e.g., *IRF7*/*9*), as well as several effector molecules involved in the regulation of inflammatory responses to infection (e.g., *IL6, TNFα* and *IL1β*).

### Stronger early innate immune response in apes than in AAM

Next, we sought to characterize differences in immune responses across species. To do so, we first looked at overall differences in the magnitude of the transcriptional responses to LPS and GARD across species (see Methods). We found that, at early time points, both ape species (human and chimpanzee) engage a much stronger transcriptional response to both stimuli as compared to rhesus and baboons (in average ∼2-fold higher, Wilcoxon test *P* < 10^−10^, **Figure 2A**). Next, we identified genes for which the magnitude of the transcriptional response to LPS or GARD was significantly different between apes and AAM (FDR<0.10 for *all* pair-wise contrasts between an ape and an AAM species and an average |log2 FC| > 0.5). Hereafter, we refer to these genes as Clade Differentially Responsive Genes, or c-DRGs. We identified a total 831 and 443 c-DRGs, in the early response (4 hours) to LPS and GARD, respectively (**Figure 2B, Table S5**). Among c-DRGs, 83-92% showed a stronger response in apes as compared to AAM, consistent with the genome-wide pattern of an overall more robust transcriptional response to immune stimulation in apes. Importantly, the stronger response observed in apes is not explained by higher baseline expression levels of the receptors involved in the recognition of LPS (*TLR4, CD14, LY96* and *CASP4*) and GARD (*TLR8*) (**Figure S2**). Next we focused our analyses on a manually curated list of 1079 genes belonging to different modules of the innate immune system^18^ and that were found to change gene expression in at least one of our experimental conditions, in at least one of the species from our dataset. These genes include sensors (n=188), adaptors (n=36), signal transducers (n=209), transcription (factors) (n=74), effector (molecules) (n=115), accessory molecule (n=54) and secondary receptors (n=50). All modules show similar divergence between clades, with ∼15% of the genes within each module classified as c-DRGs with a stronger response in apes, as compared less than 5% showing a significantly stronger response in AAM (**Figure 2C**).

**Figure 2.**
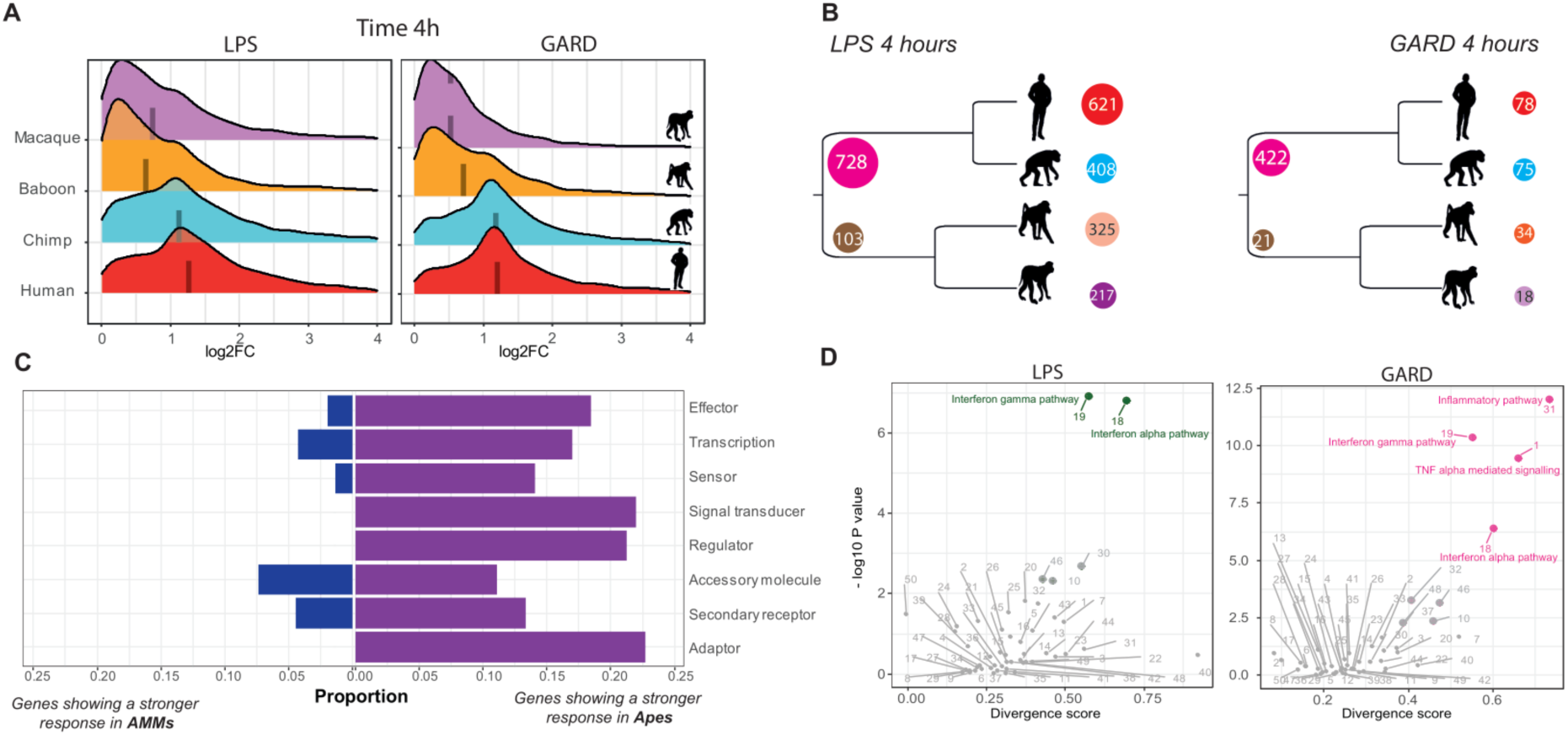
Stronger early innate immune response in apes than monkeys. **(A)** For each combination of stimulus and time-point we show the distribution of the log2 fold changes (x-axis) among genes that response to that treatment in at least one of the species. The median log2 fold change responses in each species is represented by a dashed line. **(B)** Number of differentially responsive genes that are clade- or species-specific differently regulated genes at 4 hours post LPS (left) and GARD (right) stimulation. For clade differently regulated genes (c-DRG) we report number of genes that show a stronger response in specific clade at the beginning of ancestral branch of the tree. For example, in response to LPS we identified 831 c-DRGs from which 728 show a stronger response in apes and 103 in AAMs. For species-specific responsive genes numbers are given in front of each species. The color codes for each species are red for human, cyan for chimp, orange for baboon and violet for macaque. **(C)** Bar plots represent the proportions of different classes of innate immunity genes that are classified as c-DRGs with a higher response in apes (dark violet) or in AAMs (dark blue). **(D)** Scatter plot displaying total divergence scores of hallmark pathways for LPS (green) and GARD (pink) at 4h stimulations. For a given pathway, the total divergence is given by divergence score (DS) on the x-axis and -log10 p values for each DS is on the y-axis. The pathways names, DS values, and corresponding p values are shown in **Table S6**. We highlight the pathways showing the most significant divergence scores for both the response to LPS and GARD.

To further characterize functional differences in immune regulation between apes and AAM we devised a new score of transcriptional divergence at the pathway level. We focused on the set of 50 “hallmark pathways”, which capture well-defined and curated biological states or processes.^19^ Briefly, for each gene in these pathways, a divergence score between apes and AAM was computed by calculating the average difference between the fold-change estimates between all pairs of species of the two clades, while taking into account variance in transcriptional response within each species. The pathway divergence score reflects the average divergence scores across all genes of a given pathway (see Methods for details). In the early response to LPS, the most divergent pathways between apes and AAM were “Interferon alpha response” and “Interferon gamma response” (*P ≤* 0.01, **Table S6**), indicating that the regulation of interferon responses has significantly diverged since the separation between apes and AAM. In the early response to GARD, pathways directly related to the regulation of inflammatory responses, notably TNF-*α* signaling, were the most divergent (*P ≤* 0.01) (**Figure 2D**).

### Species-specific immune responses reflect unique immune regulation mechanisms and lineage-specific divergence

Next, we sought to identify genes that respond to immune stimulation in a species-specific fashion. These were characterized as genes for which the magnitude of the response to LPS or GARD in one species was significantly different to that observed in all other species (see Methods). Across time-points and immune stimulations we identified a total of 980, 726, 425 and 655 species-specific responsive genes in human, chimpanzees, macaques and baboons, respectively (**Table S7**). Among baboon-specific responsive genes the vast majority (69%) showed a weaker magnitude of the response to 4-hours of LPS stimulation in baboons as compared to all other primates **(Table S7**). Gene ontology (GO) enrichment analyses (**Table S8**) revealed that these genes were enriched among defense response genes (FDR= 4.8×10^−14^) and a variety of other GO-associated immune terms (**Figure 3B**), including several key transcription factors (e.g., *STAT1, IRF7*/*9*), major inflammatory cytokines (e.g., *IL1B* and *CXCL8*), and a number of genes directly involved in LPS sensing and recognition (adaptor molecules *IRAK2, 3* and *4*, and the primary LPS receptor, *TLR4*) (**Figure 3A; Table S7**). The weaker response observed in baboons appears to be, at least in part, due to a higher baseline expression level of many of these innate immunity genes (**Figure S3**). Baboons have been suggested to bear higher pathogen loads than apes due to their mating promiscuity, and so it is tempting to speculate that increased baseline might represent a mechanism of protection against frequent microbial infections. ^20,21^. In rhesus macaques, the other AAM species, genes showing a stronger response to LPS at both 4 and 24 hours than that observed in all other species (N=157, **Table S7**) were mostly enriched among genes involved in the regulation of inflammatory responses (FDR = 0.002, **Table S8**), including *TREM2* a known suppressor of PI3K and NF-kappa-B signaling in response to LPS.

**Figure 3.**
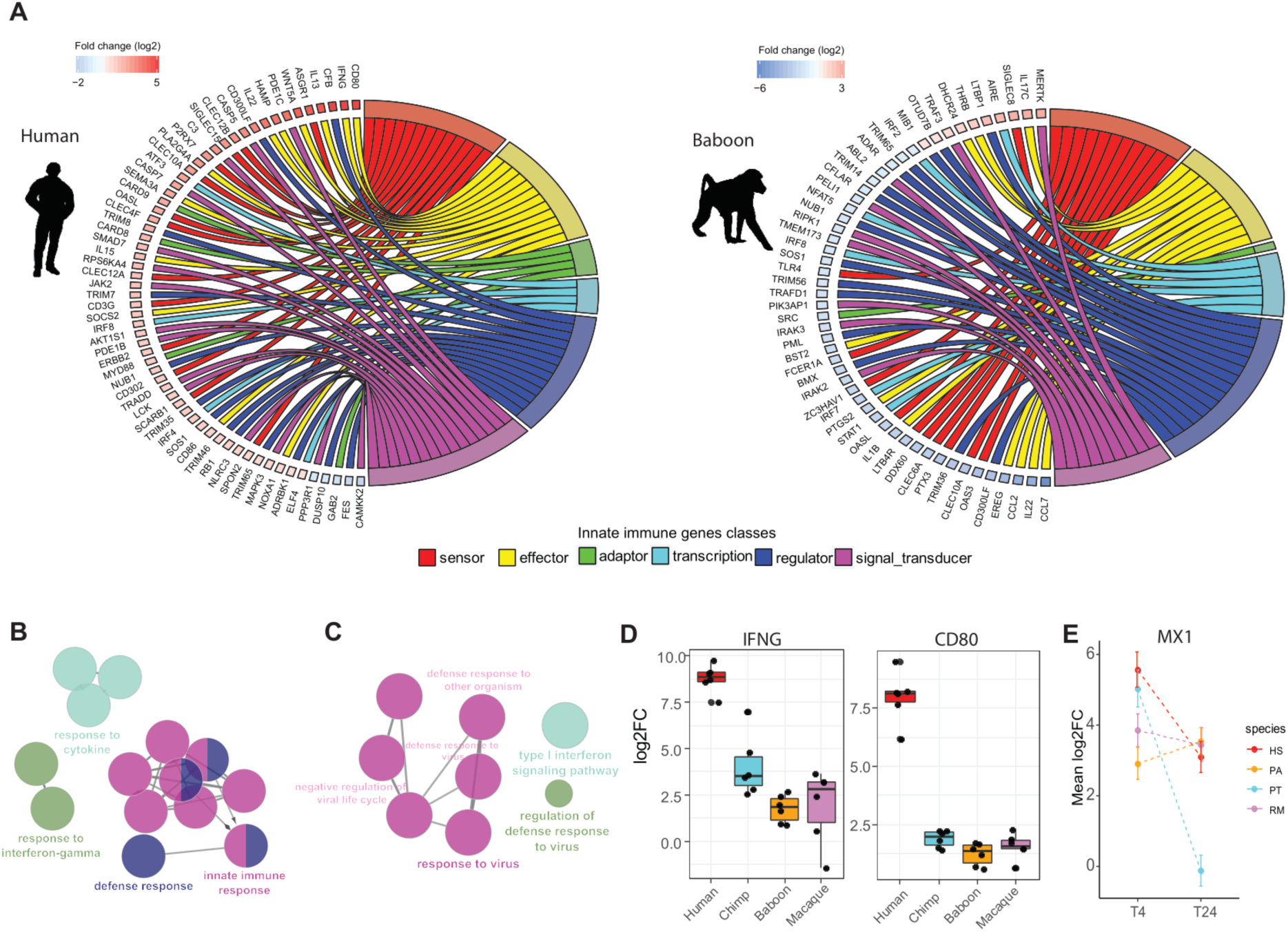
Species specific immune response reflect unique immune regulation mechanisms and lineage specific divergence. **(A)** Circos plots showing different classes of innate immune genes (clustered using different color codes) classified as species-specific responsive at 4 hours post-LPS stimulation in humans (left) and baboons (right). The log2FC key represent the average difference between species response versus all other species where the positive (red color) and negative (blue) values indicate the magnitude of the stronger and weaker absolute response in this species vs. all others, respectively. (**B**) Gene ontology (GO) enrichment analysis for genes showing an weaker response to LPS at 4 hours in baboons as compared to all other species. **(C)** Gene ontology (GO) enrichment analysis for genes showing an weaker response to LPS at 24 hours in chimpanzees as compared to all other species. For B and C only top GO terms are presented. The full list of significant GO terms can be found in **Table S8. (D)** Boxplot represent the log2FC of *IFNγ* and *CD80* genes which, at 4 hours post-LPS stimulation, were found to have a significantly stronger response in human than in other primates. (E) Estimates of the mean fold changes response for *MX1* (+/- SE) at the two time points across the four primate species studied.

Among chimpanzees-specific genes the most notable GO enrichments were observed among genes showing a weaker response to LPS at 24 hours relative that observed in all other primates. These genes were significantly enriched for GO terms associated with viral defense mechanisms, including “response to virus”, or “type I interferon signaling pathway” (FDR<1×10^−9^, **Figure 3C**). Further inspection of these genes revealed that the vast majority are strongly up-regulated at 4 hours post-LPS stimulation – at similar levels to those observed in other species - but that chimpanzees have a unique ability to shutdown these genes at later time points. For example, the prototypic interferon responsive gene *MX1* is up-regulated by over 5-fold in all primates at 4 hours but by 24 hours *MX1* levels have revert to baseline uniquely in chimpanzees (**Figure 3E**), suggesting that chimpanzees are particularly divergent in the regulatory circuits associated with the control of viral responsive genes.

In contrast to the pattern observed for baboons, human-specific responses were associated with genes showing a stronger response to immune stimulation as compared to that observed in other primates. Gene ontology analyses revealed that these genes are over-represented among terms related to the regulation of cytokine production involved in immune response (FDR = 0.045), and T cell activation involved in immune response (FDR = 0.06) (**Table S8**). Notable examples of human-specific responding genes include the canonical T cell co-stimulatory molecule CD80 (average 5-fold increase in response to both stimuli relative to other species) and *IFNγ*, a cytokine central for protective immunity against a large number of infectious agents and the key determinant of the polarization of T cells towards a pro-inflammatory Th1 phenotype^22^ (**Figure 3D**). The higher production of *IFNγ* and *CD80* in humans may mediate more effective killing of viral and bacterial pathogens. Further, as these molecules are important regulators of cytokine production and T cell activation, it also suggests significantly different regulation of T cell responses.^23,24^

### Regulatory divergence decreases as infection proceeds

Next, we compared the transcriptional divergence between early (4 hour) and late (24 hours) immune responses. We observed a marked reduction in divergence scores at 24 hours post-stimulation of most hallmark pathways in the response to both LPS (*P*=8×10^−6^) and GARD (*P*=6×10^−9^) (**Figure 4A**). In LPS-stimulated cells, the most striking differences were observed for interferon-related pathways, which show a reduction in divergence score of ∼6-fold between the two time points. In GARD-simulated cells, the largest reduction in divergence scores was observed among pathways related to the regulation of inflammatory responses (**Figure 4A**). These findings indicate that most transcriptional divergence in immune responses among primates occurs during the initial response to pathogens followed by an overall convergence to similar response levels at later time point, specifically among genes involved in the regulation of inflammation and viral-associated interferon responses (**Figure 4B**). In apes (but not in AAMs), genes involved in the regulation of inflammation are strongly enriched among those for which the response to GARD significantly decreases at the later time point, whereas those decreasing in response to LPS are enriched for viral response genes (**Figure 4C**; **Table S9**).

**Figure 4.**
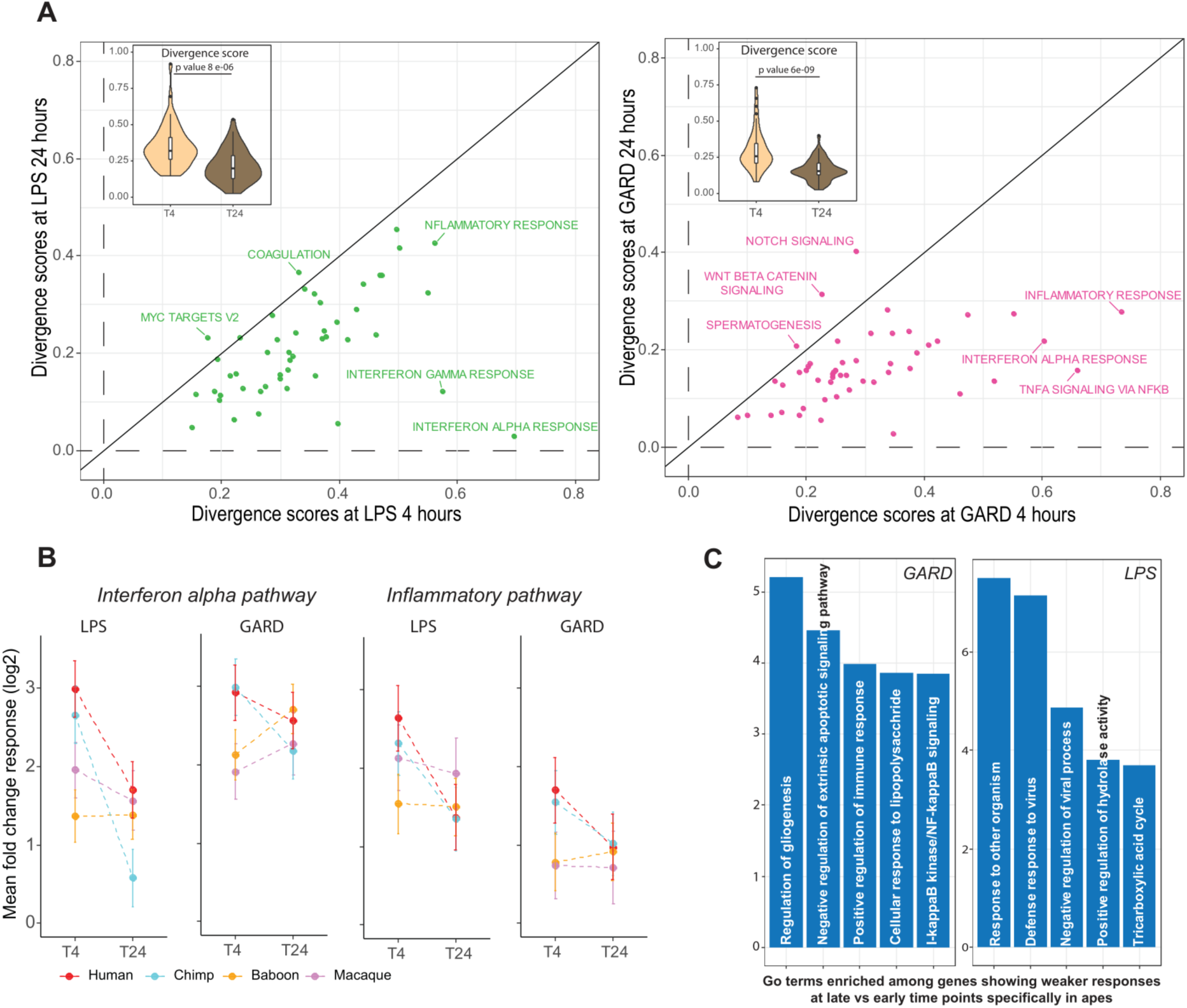
Divergence of immune response is reduced at later time point. **(A)** Scatter plots of divergence scores of hallmark pathways at early (x-axis) and later time points (y-axis) for LPS (green) and GARD (maroon) stimulations. The inset boxplots contrast the distribution of divergence scores among all pathways between the two time points. P values were obtained using Mann Whitney test. **(B)** Estimates of the mean response at the two time points for each species (+/- SE) across genes bellowing to the interferon alpha and inflammatory response hallmark pathways. **(C)** GO enrichment analysis for genes that showed significant decrease in response in apes only (FDR < 0.05 in apes and FDR > 0.05 in monkeys) for LPS and GARD. Top significant GO terms are given as indicated by –log10 p value on the x axis.

### Apes engage a less specific innate immune response than AAM

An aspect of innate immunity central to its success during microbial assault is its ability to recognize pathogens and initiate the most appropriate defense against them by type. The specificity of the innate immune response to infection is mediated by pattern recognition receptors that detect the presence of danger signals via conserved molecular patterns associated with subtypes of pathogens and host damage (e.g. penta- and hexa-acylated LPS from Gram-negative bacteria detected by *TLR4*-*LY96* receptors).^25,26^ Signals of viral danger such as GARD are expected to activate a response mainly controlled by transcription factors prominent in antiviral defense such as interferon regulatory factors (IRFs), which limit viral replication and dissemination through the upregulation of interferons and interferon-regulated genes.^27,28^ By contrast, recognition of Gram-negative bacteria via cell-wall component LPS stimulates a broader array of cytokine responses that tends towards expression of pro-inflammatory cytokines regulated by transcription factors *NFκB* and *AP1*, but can also include interferon expression regulated by transcription factors such as *IRF3* and JAK-STAT.^27 29-31^

Two major lines of evidence indicate that early transcriptional immune responses are less specific in apes than in AAM. First, we found the transcriptional responses to LPS and GARD were more similar to each other in apes (humans r = 0.87, chimpanzees r=0.83) than they were in baboons (r = 0.44) or macaques (r = 0.65) (**Figure 5A**). Accordingly, we found about three times more genes that respond *uniquely* to either LPS or GARD (i.e., “ligand specific” genes) in AAM as compared to apes (chi^2^ test; P=2.2×10^−16^)(**Figure 5B**, Methods for details on the statistical model used to characterize ligand specific and shared genes). The second piece of evidence comes from the nature of the genes that are differentially activated in response to LPS and GARD. The fact that apes show a higher correlation in GARD and LPS responses compared to AAMs predicts that they will tend to activate both antibacterial and antiviral defenses mechanisms regardless of the nature of the stimuli. Supporting this notion, the genes that exhibited a stronger response in apes than in AAMs after stimulation with the viral mimic GARD for four hours were most significantly enriched genes involved in the regulation of inflammatory responses (*P*= 7.1×10^−6^; FDR = 0.008, **Table S8**), whereas genes involved in the response to viruses (GO term “response to virus”) were enriched upon bacterial LPS stimulation (*P*= 0.0023; FDR = 0.15, **Table S8)**.

**Figure 5.**
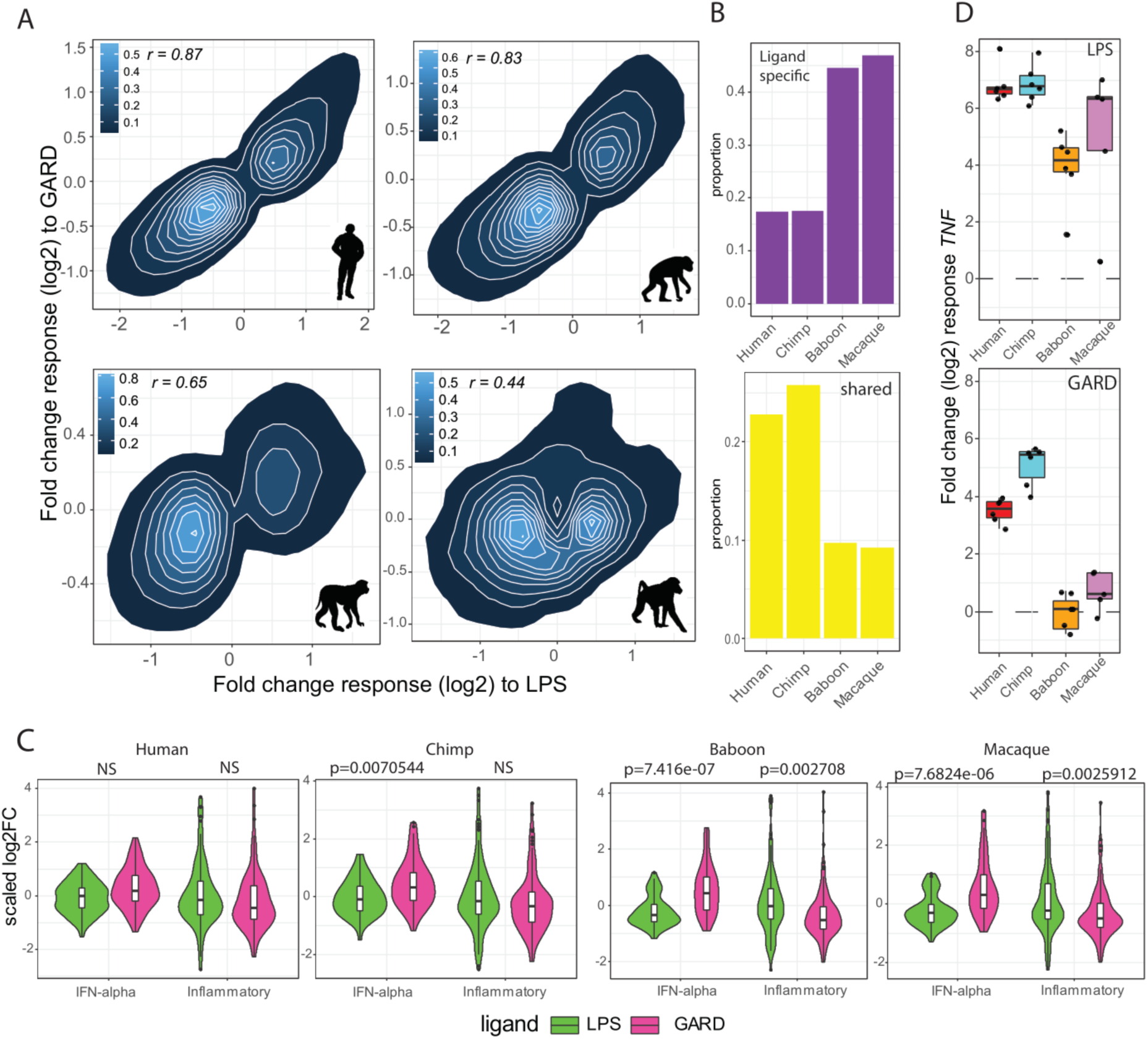
Apes engage a less specific innate immune response than AAMs. **(A)** Correlation plots of the magnitude of the fold change responses between LPS (x-axis) and GARD-stimulated cells (y-axis). For each of the species, we only include genes that were differentially expressed (FDR < 0.05) in response to at least one of the stimuli (N= 7862, 7874, 6585 and 5430 genes for human, chimp, baboon and macaque, respectively). High correlation was found in apes (∼ 0.85) while modest correlation was found in baboon (0.44) and moderate in macaque (0.65). **(B)** Proportion of ligand-specific (i.e., genes that respond uniquely to either bacterial or viral stimuli) and shared genes (i.e., genes equally activated by both immune stimuli) across species. **(C)** Violin plots comparing (scaled) log2 fold-change responses to 4 hours of LPS and GARD stimulation between genes belonging the hallmark pathways “Interferon (IFN) alpha” and “inflammatory response”. The p-values shown have been Bonferroni corrected for the number of tests performed. “NS” stands for non-significant (i.e., p value > 0.05) **(D)** Boxplots of the log2 fold change response (y-axis) of *TNF* in response to LPS and GARD stimulations across primates.

To explore these differences in more detail, we focused on genes involved in the interferon alpha pathway (viral-associated response) or inflammatory response (bacterial-associated response). In AAMs, inflammatory response genes tended to be more strongly up-regulated in response to LPS compared to GARD (*P*≤ 0.0027), suggesting that their transcription is particularly sensitive to receipt of a bacterial danger signal compared to a viral one. No significant differences in upregulation of these same genes were noted between LPS and GARD cells in apes () (**Figure 5C**). For example the canonical pro-inflammatory cytokine *TNF*, which in macaques and baboons is strongly up-regulated only in response to LPS, is potently up-regulated in response to both stimuli in humans and chimpanzees (by over ∼4-fold, **Figure 5D**). Other examples of this pattern include the classical pro-inflammatory cytokines *IL1A* and *IL1B* (**Figure S4**). Likewise, interferon-associated genes were more strongly up-regulated in response to GARD compared to LPS in AAM (P<=7.7×10^−6^), while in apes these genes showed more concordant levels of up-regulation between stimuli (**Figure 5C**). Interferon-induced and potent antiviral genes, including *MX1* and *OAS1*, were much more strongly upregulated in response to GARD than to LPS in AAMs compared to apes (**Figure S4**).

## Discussion

Our study provides a genome-wide functional comparison of variation in innate immune responses between species belonging to two closely related clades of primates. Ape (human and chimpanzee) total blood leukocytes were significantly more responsive to bacterial and viral stimulation compared to total blood leukocytes obtained from AAM (rhesus macaques and baboons) during the early hours of challenge, mounting generally stronger and less specific transcriptional responses. This increased response suggests apes maintain increased sensitivity to particular types of microbial assaults compared to AAM, a phenomenon likely to come with considerable energetic cost.^1,5^ From an evolutionary standpoint investment in increased sensitivity to pathogens can limit the negative effects of pathogen exposure on reproductive fitness. Humans and chimpanzees participate in a comparatively slower life history than rhesus macaque and olive baboon monkeys – they live decades longer, take longer to reach sexual maturity, nurture their young longer and maintain a larger body size^32-34^. A long life at a large size increases risk of pathogen exposure both in terms of number of exposures and absolute load, over the course of a life that will have long periods of time between the birth of offspring. A slow life history strategy can be concomitant with and increase risk in pathogen-mediated limitations in reproductive success, making a more substantial investment in robust early pathogen detection and elimination evolutionarily beneficial, compared to the ordinary metabolic costs of launching those responses.^35,36^

However, serious bystander tissue damage is a cost for immune protection during severe infections. Pathogen virulence may play a significant role in the evolution of high energy low specificity early immune responses. The primate genera in this study substantially differ in their evolutionary exposure to particular pathogens (e.g. dengue virus, immunodeficiency viruses, Zika virus)^37-40^. Exposure to pathogens of high virulence may lead to a low cost-benefit ratio for primate hosts, since the reproductive and evolutionary benefit of a transiently demanding immune response outweighs its energetic and tissue costs. ^41,42^. Under this rubric, a robust but less specific early response to pathogens is effective and beneficial most of the time. Any contribution that response might make to immunopathology in apes through potentially increased risk of sepsis or chronic inflammatory disease is evolutionarily negligible compared to the persistent risk of infection. Interestingly, among the most divergently responding pathways between apes and AAMs, several were associated with the regulation of interferon responses and responses to viruses. These findings are consistent with growing body of literature that pathogens and, specifically, viruses have been important drivers of adaptive evolution in humans and other mammals.^15,43-45^

Regardless of initial strength and divergence of transcriptional response to LPS and GARD, we show that the transcriptional activity of antiviral (interferon) and inflammatory pathways became attenuated over time and more similar between species. While acute-phase and early proinflammatory responses are typically later countered by a later anti-inflammatory response to lessen host damage and maintain homeostasis, the dampening of this initial powerful antimicrobial response over time, is profound^46^. Remarkably, in apes the pathways that underwent the most pronounced attenuation after 24h tended to be ones not expected to be strongly engaged in the response to the pathogen type in the experiment. For instance, the typically antiviral type I IFN pathway response was found to be markedly reduced in apes after 24 hours of bacterial but not viral stimulation. While the initial response of apes to immune stimulus is very strong, temporal regulation of responding pathways may reduce the energetic costs of such an immune strategy. What gene regulatory and immunological mechanisms are involved in such temporal regulation will require further investigation.

In conclusion, we show initial antibacterial and antiviral responses of apes to be highly correlated, and strongly responsive when compared to close relatives African and Asian monkeys. Apes appear to have adopted an immune strategy that emphasizes sterilization over specificity, strongly transcribing a greater number of genes in response to immune stimulation and releasing very similar immune transcriptomic “arsenals” regardless of pathogen-type. This powerful response dramatically shifts during the opening hours of infection, to involve significantly fewer genes after 24 hours, which may help limit bystander tissue damage. The energetically costly approach apes initiate in response to immune stimulation may be favored by this primate family’s adoption of slower life history with increased risk of pathogen exposure over reproductive life span, or past pathogen exposure. The addition of more primate species, combined with the use of single-cell RNA sequencing methods are important next steps to study the evolution of the immune system and more precisely map the immune cell types that contribute the most to divergence in immune response across primates.

## Materials and methods

### Sample collection and blood stimulation

We measured innate immune responses on a panel of 6 humans, 6 chimpanzees, 6 rhesus macaques, and 8 olive baboons (three females and 3 males for each species, 4 females, 4 males for baboon). Human samples were obtained via informed consent, with the approval of the Research Ethics Board at the Centre Hospitalier Universitaire Sainte-Justine (Research Ethics Board approved protocol #3557). Non-human primate blood samples were humanely collected in accordance with the animal subject regulatory standards of the Texas Biomedical Research Institute and Emory University Institutional Animal Care and Use Committees. Chimpanzee samples were collected prior to the NIH ban on chimpanzee research.

We drew 1 mL of whole blood from each animal directly into a TruCulture tube (Myriad RBM) that contained: (i) cell culture media only (“control”), (ii) cell culture media plus 1 µg/mL ultra-pure LPS from the *E. coli* 0111:B4 strain (“LPS”),or (iii) cell culture media plus 1 µg/mL of Gardiquimod (“GARD”). Samples were incubated for 4 and 24 hours at 37°C. Following incubation, we separated the plasma and cellular fractions centrifugation, and lysed and discarded the red cells from the remaining cell pellet by applying red blood cell lysis buffer (RBC lysis solution, 5 Prime Inc.) for 10 minutes followed by centrifugation and washing with 1x PBS. The remaining white blood cells were lysed in Qiazol and frozen at -80C until library construction (Qiagen, San Diego, CA, USA). To control for variation in cellular composition in downstream analyses, we used flow cytometry to quantify the proportions of leukocyte subtypes, accounting for polymorphonuclear (CD14dim/SSC-A>100K/FSC-A>100K/CD66+), classical monocytes (CD14+/CD16-), CD14+ intermediate monocytes (CD14+/CD16+), CD14- non- classical monocytes (CD14-/CD16+), helper T cells (CD3+/CD4+), cytotoxic T cells (CD3+/CD8+), double positive T cells (CD3+/CD4+/CD8+), CD8-B cells (CD3-/CD20+/CD8-), CD8+ B cells (CD3-/CD20+/CD8+), natural killer T lymphocytes (CD3+/CD16+), and natural killer cells (for monkeys: CD3-/CD16+ in the lymphocyte scatter, for apes: CD3-/CD16+/CD56+ in the lymphocyte scatter) Samples for FACS were simultaneously cleared of red blood cells vis lysis and fixed by application of BD FACS-lyse for 2 minutes, prior to washing with 1x PBS, staining with fluorochrome conjugated monoclonal antibodies (**Table S10**), before washing with 1x PBS and suspending in a 1x PBS and paraformadelhyde solution for analysis on the BD LSRFortessa platforms. Proportional analysis was completed in FlowJo X, using BD FACSBeads individually stained with the antibodies to calculate compensation.

### RNA-seq data generation

#### Library construction

Total RNA was isolated from cell lysate by phenol::chloroform extraction and spin-column (miRNAeasy kit, Qiagen, San Diego, CA, USA), quantified by spectrophotometry and assessed for quality using the Agilent 2100 bioanalyzer (Agilent Technologies, Palo Alta, CA). Samples with no evidence of RNA degradation (Integrity number >8) were then used for RNA library development. Messenger RNA (mRNA) was isolated by magnetic bead and converted into RNA libraries using the Illumina TruSeq RNA Library preparation kit v2 according to the manufacturer’s instructions (Illumina, San Diego, CA, USA). Libraries were sequenced on a HiSeq 2100, producing 151 transcriptomes, at 25-30 million reads per sample

### Reads mapping on 1:1 orthologs

Following sequencing, we trimmed Illumina adapter sequence from the ends of the reads and remove bases with quality scores < 20 using Trim Galore (v0.2.7). We used STAR to align the reads to an orthologous reference genome for all four species^47^. We developed this genome using the XSAnno pipeline which combines whole genome alignment, local alignment and multiple filters to remove regions with difference in mappability between species^48^. XSAnno pipeline identifies orthologous genes across two species using three major filters namely LiftOver to carry annotation of one species over the other, BLAT aligner to compare orthologous exons identity between the two species and simNGS to identify exons that have different lengths between the species. We used the genome assemblies of hg19 for human, CHIMP2.1.4 for chimpanzee, MMUL 1.0 for macaque and PapAnu2.0 for olive baboon species. We used human annotation as a reference. The pairwise alignment chains between human and each species were obtained from UCSC genome browser.^49^ We used different thresholds to define orthologous regions between the two genomes to carry annotation from one species to another using AnnoConvert program that utilize LiftOver according to simulations using liftOverBlockSim PERL script from XSAnno pipeline^50^. The values were 0.98, 0.92 and 0.91 for chimp, baboon and macaque respectively that were used to assign - minMatch argument in AnnoConvert. Second step is using reciprocal whole genome alignment using BLAT through BlatFilter software of the pipeline using annotations files generated previously^51^. This step will filter exons that are highly divergent between the two species. The last filter is using simNGS to simulate reads for exons assuming they are not differentially expressed. Then, differential expression analysis is performed and if exons were found to be differentially expressed, these will be filtered out as it reflects differential length of the exons between species.

Gene expression estimates were obtained by summing the number of reads that mapped uniquely to each species annotated genome using HTSeq-count (v0.6.1)^52^. After excluding samples that did not produce sequenceable libraries and post-sequencing quality control, we analyzed read counts for 151 samples (Humans: 12 controls, 12 LPS, 12 GARD; Chimpanzee: 12 controls, 12 LPS, 12 GARD; Rhesus: 11 controls, 12 LPS, 11 GARD; Baboons: 16 controls, 14 LPS, 15 GARD; **Table S1**). We confirmed the identity of all samples based on genotype information derived from SNP calls made from the RNA-seq reads.

### Read normalization and filtering lowly expressed genes

Prior to RNA-seq data analysis, we first filtered out genes that were very lowly or not detectably expressed in our samples. Specifically, we only kept genes whose expression was higher or equal to one count per million (CPM) in all the individuals from at least one species, and one of the experimental conditions. This procedure yielded a total of 12,441 genes used for further analysis. Normalization for sequencing depth and library sizes was done using Trimmed Mean of M-values (TMM) method.^53^ We normalized the resulting read count matrix using the function *voom* from the R package limma to allow using linear models by limma package.^54^ The voom algorithm models mean-variance trend of logCPM for each gene and uses it to compute the variance as a weight of logCPM values. We then modeled the normalized expression values as a function of the different experimental factors in the study design such as species, ligand and time points.

### Statistical analysis

All statistical analysis was done on R version 3.6.2. Differential expression analysis was done using limma package v.3.34.9.^55^ We employed linear regression to identify DEGs according to different questions asked by designing different models. We designated a model to test for differences of gene expression across species and treatments, ∼ covariates + species + species:Time.point.stimulant. The arm Time.point.stimulant is the samples for each experimental condition i.e. LPS.4h, LPS.24h, GARD.4h and GARD.24h. From this design, one can retrieve ligand responses in each species right away, while responses to ligands at 24h are built from linear combinations, such us (LPS.24h -NC.24h) and (GARD.24h -NC.24h). To take into account the paired structure of the data, with different samples coming from the same individuals, we used the duplicateCorrelation function. The used covariates are the different cell proportions collected by the FACS data. The cell proportions covariates are aimed to correct for the different proportion of white blood cells in different primate species since we conducted the transcriptomic characterization on all immune white blood cells. Genes with different magnitude of response between clades, referred to as clade differentially responsive (c-DR) genes were characterized. We established two filters to characterize significant c-DR genes in each treatment. Firstly, we required the genes not to be differentially responsive to the treatment, even marginally, between within clade species pairs (chimp vs human and baboon vs macaque showing FDR>0.25). Second, we required that any pairwise comparison involving species from different clades to be significant at FDR<0.1. Third, we also computed the average differences in reponses between apes and AAMs, as follows: the absolute difference between the average response in apes vs AAMs, as follows:

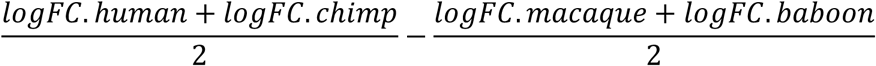

And required that contrast to be significant at FDR<0.1, with genes featuring

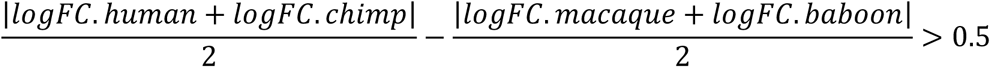

being labeled ape-specific; and AAM-specific for those for which:

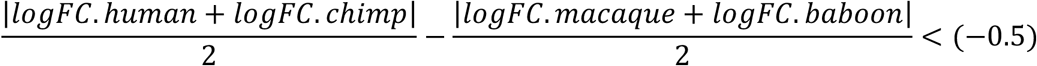

Species-specific differentially responsive (s-DR) genes were identified using pairwise comparisons at FDR < 0.01; consistent direction of expression in all contrasts, and systematic differences corresponding to stronger, or weaker responses in the species of interest with respect to the any of the other three. Finally, we also required genes to show a logFC in response to the stimulus whose absolute differs in more than 1 log2FC with respect to the average of the other three animals. For humans, as an example, this means that:

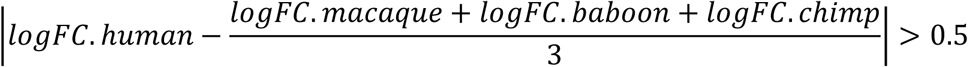

Ligand specific genes in each species are genes that are respond to one ligand (FDR<0.05), but not to the other (FDR>0.25); and whose responses to both ligands are in turn significantly different (FDR<0.05). Shared genes are those whose responses to ligands are both significant (FDR<0.05 in both), and, at the same time, not significantly different between them (FDR>0.25) Correction of multiple testing was done using false discovery rate, FDR, as described by Benjamini-Hochberg.^56^

### Divergence scores

For each time-point and stimulus, species were compared pairwise to retrieve the absolute differences between species’ responses to the stimulus under analysis. For the pair chimp vs human, for example, we can define:

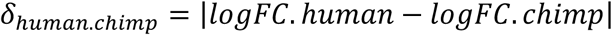

Comparing these differences for pairs of animals within versus across clades, we obtained divergence scores as follows:

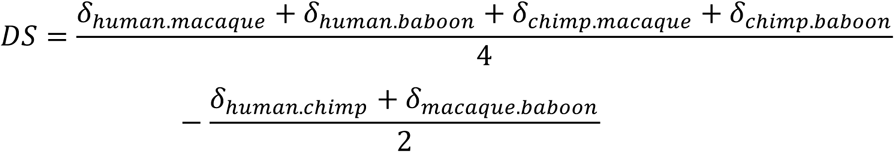

The analysis was conducted for all 50 hallmark pathways. We restrict the analysis in a given pathway to responsive genes (FDR < 0.05 in any species), whose average DS is reported. A p value for each DS of a given pathway was calculated by contrasting the DSs of genes of this specific pathway against the DSs of all responsive genes using Wilcoxon test.

### Functional characterization

We conducted the functional characterization using gene ontology (GO) enrichment implemented in CluGO application (2.5.5) of Cytoscape (v.3.7.2) ^57^ Benjamini-Hochberg method for multiple correction was used and all orthologous genes, 12441 genes, were used as a background. Default values were used for the rest of the parameters. FDR cut off use was below 0.15.

### Data Availabilty

The RNA-seq data generated in this study have been deposited in Gene Expression Omnibus (accession number XXX).

## Author Contributions

L.B.B. and J.F.B. designed research, M.B.F.H., J.F.B., J.K., J.S., J.C.G., Y.V., and L.B.B. performed the research, M.B.F.H., J.C.G., J.F.B. and L.B.B. analyzed the data, M.B.F.H., J.F.B. and L.B.B. wrote the paper with contributions from all authors.

## Acknowledgements

The authors thank Steven Bosinger, and Guido Silverstri of Yerkes Primate Center and Emory University for their assistance acquiring samples. We thank L.B.B. laboratory members for critical reading of the manuscript. We thank Calcul Québec and Compute Canada for providing access to the supercomputer Briaree from the University of Montreal. This work was supported by RGPIN/435917-2013 from the Natural Sciences and Engineering Research Council of Canada (NSERC) and R01-GM134376 from the National Institute of General Medical Sciences to L.B.B.. JFB is funded by NSF-BCS-1750675. The resources of the Southwest and Yerkes National Primate Research Centers are supported by NIH grants P51-OD011133 and P51-OD011132, respectively, from the Office of Research Infrastructure Programs/Office of the Director

## Conflicts of Interest

The authors have no conflicts of interest

## Supplementary Tables

**Table S1:** Summary of all samples included in this study and associated metadata.

**Table S2:** Pairwise comparisons of cell population proportions between primate species.

**Table S3:** Summary of log2FC and FDR for all orthologs for all species across all treatments.

**Table S4:** Conserved genes that are responding in all species in LPS and GARD 4h treatments.

**Table S5:** Clade differentially responsive genes across all treatments.

**Table S6:** Divergence scores and p values for all hallmark pathways in all treatments.

**Table S7:** Species-specific differentially responsive genes. Two sets are provided for every species, genes with higher response in this species vs. all other species (species.DRSG.higherRes) and genes with lowest response compared to all other species (species.DRSG.lowerRes).

**Table S8:** Gene ontology enrichment analysis of all clade and species specific differentially responsive genes.

**Table S9:** List of genes underwent temporal reduction or increase in response at later time point in each species and treatment (LPS and GARD).

**Table S10:** List of antibodies used to characterize primate’s leukocytes composition.

## Notes

### Competing Interest Statement

The authors have declared no competing interest.

