## Supplementary Figures for "Primate innate immune responses to bacterial and viral pathogens reveals an evolutionary trade-off between strength and specificity"

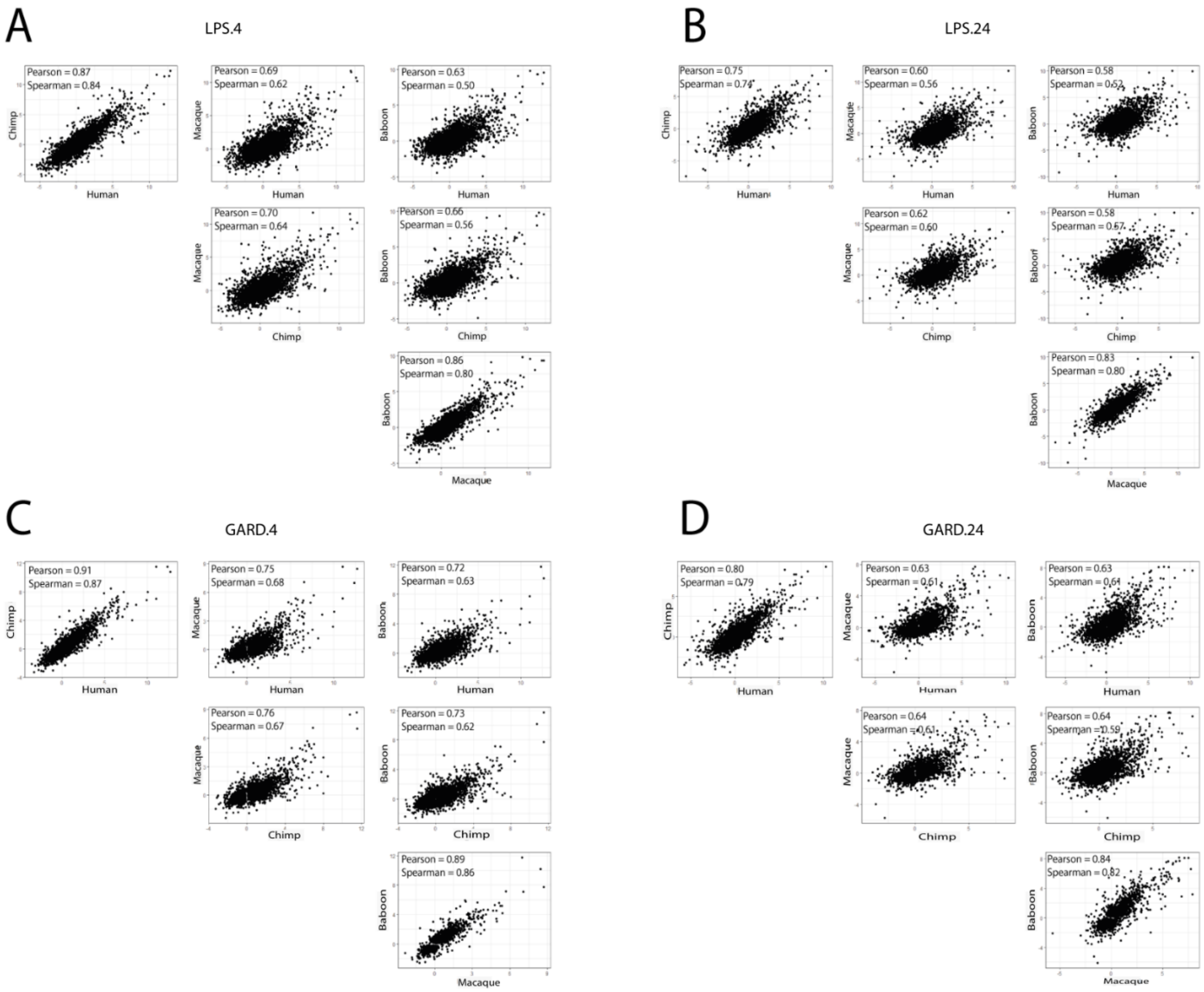


**Supplementary figure S1**: Correlation across species between fold-change (log2 scale) responses among genes that are either significant in LPS treatment at 4 and 24h (A and B respectively) or GARD treatment at 4 and 24h (C and D respectively) at FDR < 0.05 in at least one of the species studied. Spearman and Pearson correlation coefficients are indicated on each plot.

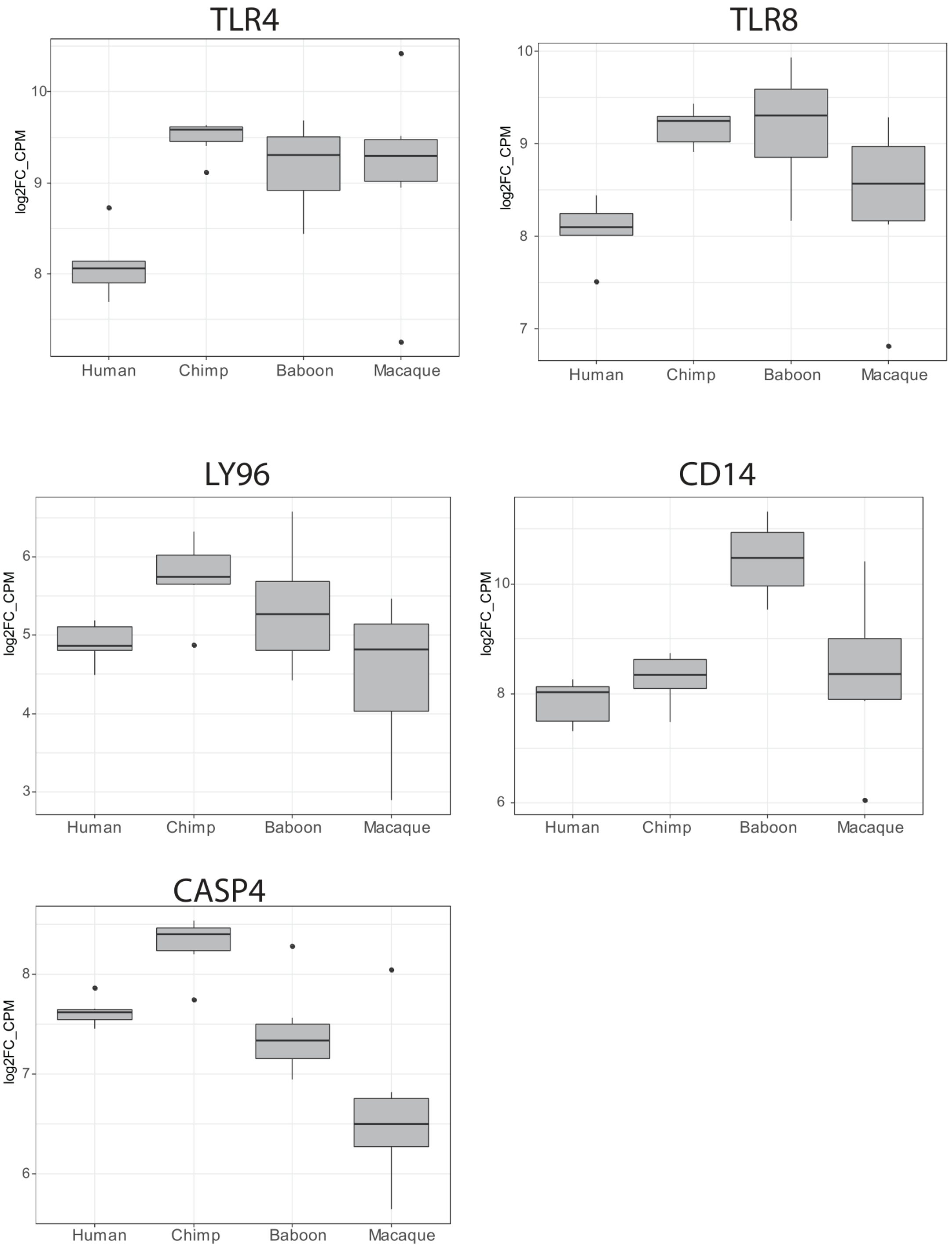


**Supplementary figure S2**: Boxplots representing expression values across species (log2 count per million; log2CPM) of key sensors involved in the recognition of the ligands used in this study, LPS and GARD.

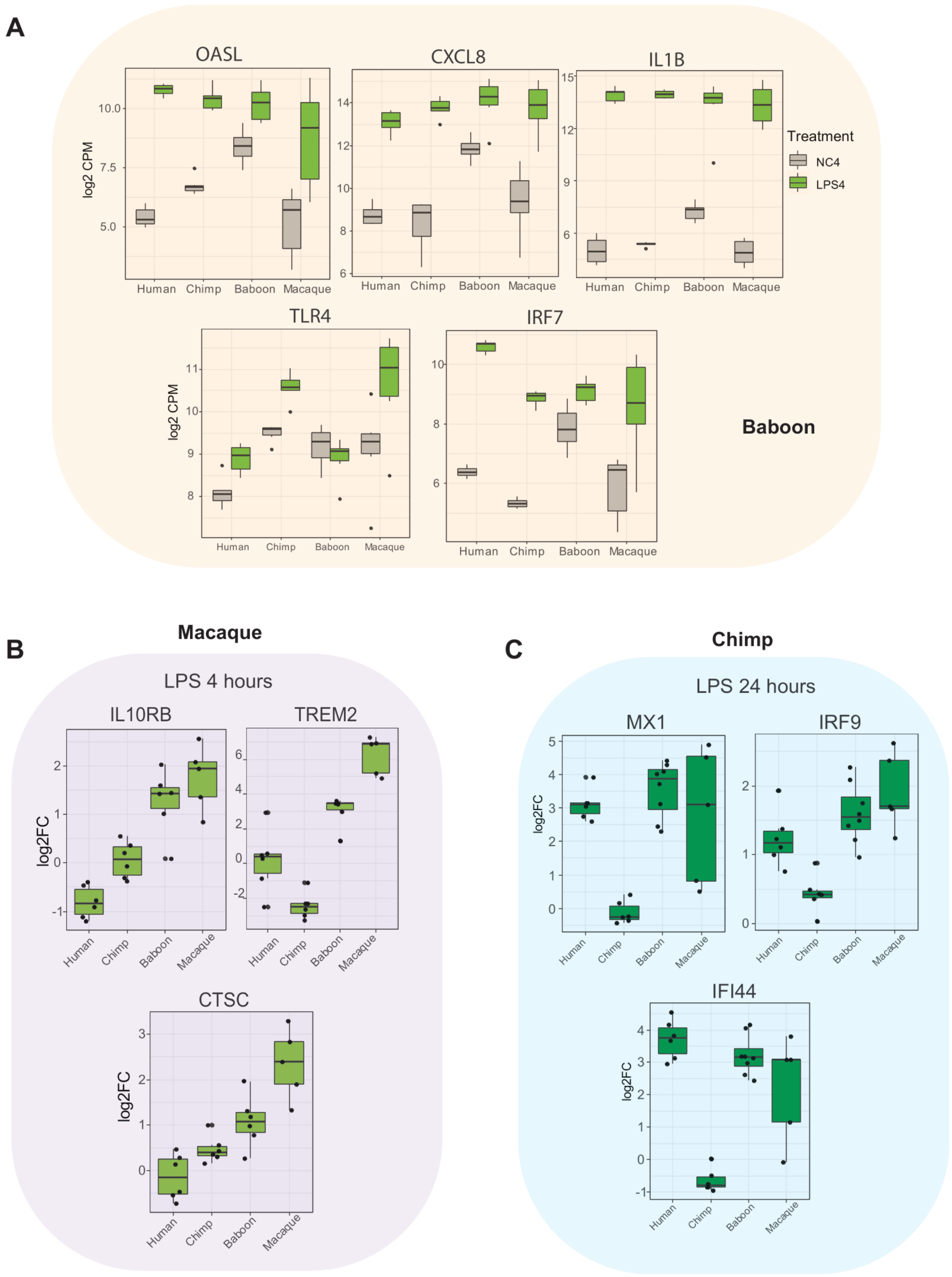


**Supplementary figure S3**: **(A)** Example of baboon-specific genes. The boxplots represent the expression values across species in non-stimulated cells (gray) and cells stimulated with LPS for 4 hours (green). In all cases showed herein baboons show a weaker response to LPS compared to all other species, which is primarily due to an increased baseline expression of these genes. **(B)** Example of rhesus-specific genes at 4 hours post LPS stimulation. **(C)** Example of chimpanzee-specific genes at 24 hours post LPS stimulation.

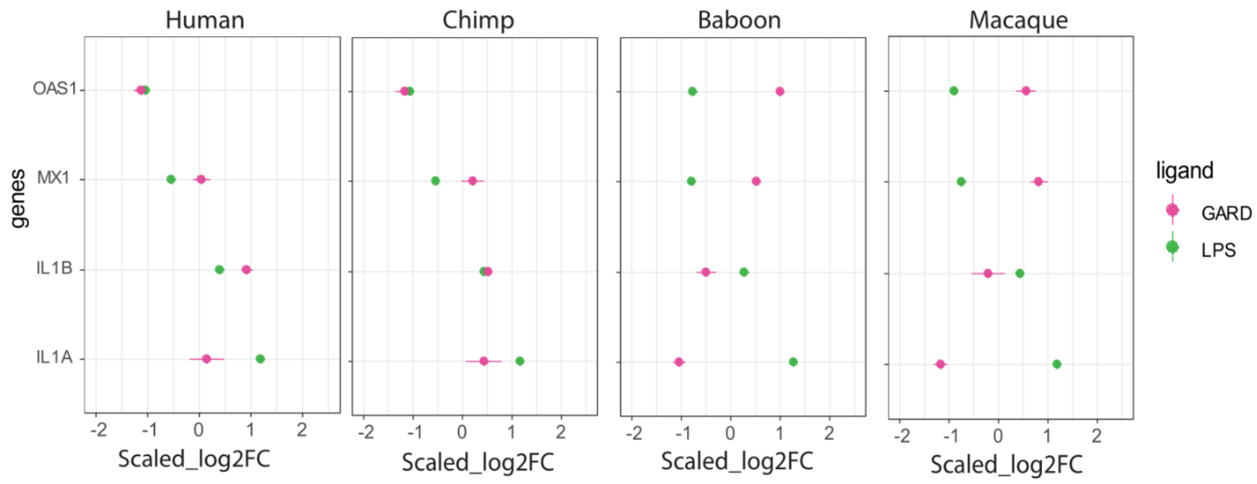


**Supplementary figure S4** Scaled log2FC of number of key innate immune genes that showed distinct response to bacterial or viral ligands in monkeys vs apes at 4h time point.
